# Predicting Fixation Paths in the Moran Process:A Machine Learning Approach

**DOI:** 10.1101/2023.07.14.549103

**Authors:** Mahdi Hajihashemi

## Abstract

Path of Fixation in evolutionary process highly depends on structure of underlying population. In this paper, we apply a machine learning method to predict the path of fixation in several complex graphs and two regular graphs. In our approach, the path of fixation is not used as the target variable in the machine learning model. Rather, we focus on predicting the probability of progression forward (referred to as *λ* in the literature) using the machine learning model. By using previous achievements in determining the fixation path for the Moran process, obtaining the path of fixation becomes straightforward. Due to the time and computational resources required for simulating an evolutionary process in a large population, utilizing a machine learning method can help us save both of these valuable resources. This approach can provide insights to researchers studying evolutionary processes in the context of meta-population problems.

## 1 Introduction

Evolutionary graph theory (EGT) is the most well known method for studying evolution of individual populations [1–13]. In many cases the population is not simple and we need a tool for studying diffusion like process in populations [14–19]. Graphs are considered the most effective tools for this purpose. In EGT, individuals within populations are represented as nodes, while the relationships between them are depicted through edges. Therefore, the structure of populations is summarized by the topology of graphs [20–31].

In EGT, the dynamics of evolution are governed by the Moran process. In most cases, the interactions between two or more individuals are considered and studied. In this paper, we focus in two types individuals namely residents and mutants. In Moran process the variable that indicates the power of reproduction is fitness. Fitness of residents considered 1 and fitness of mutants considered *r >* 1. The update rule is as follows: in each step of the process, one individual is selected for reproduction. The probability of selection for each individual is proportional to its fitness. The selected individual then chooses one of its neighbors in the population and replaces it with its offspring. Consequently, in each time step, the number of mutants could increase or decrease by one or remain unchanged.

In most cases, the dynamics begin with a single mutant in the population, while all other members of the population are residents. In other words, in a graph with a size of *N*, there is only one node labeled as a mutant, and all the remaining *N*−1 nodes are labeled as residents. In these circumstances, the system has two possible outcomes. The first is when the single mutant reproduces and occupies the entire population, resulting in mutants fixing the population. The second outcome occurs when residents reproduce and suppress the single mutant, leading to residents fixing the population. This update rule introduces two important concepts in the Moran process [32–37]. The first concept is the fixation probability, which refers to the probability that a single mutant will fix in the population. The second concept is the conditional fixation time, which represents the average number of steps it takes for a single mutant to fix in the population.

Suppose the single mutant reproduces and the number of mutants reaches *j*. In this situation, the probability of the number of mutants increasing to *j* + 1 is denoted by *λ*_*j*_ while the probability of the number of mutants decreasing to *j* − 1 is denoted by *μ*_*j*_. Therefore the probability that the number of mutants remains unchanged is represented by 1−*λ*_*j*−_μ_j_.

Although fixation probability and fixation time are important concepts, they do not provide us a clear picture of the evolutionary path. By “evolutionary path” we mean given that the process ends in fixation of mutants, in each step of time how many mutants are in the population. An effective tool for analyzing diffusion-like process such as Moran process is Markov chain [38–40]. In reference [41], a method is introduced that establishes a duality between the dynamics of a population in the Moran process and the dynamics of a Markov chain. This method allows for obtaining the path to fixation by knowing the value of *λ*_*i*_ for all values of *i*. Using this method by having size of population (*N*), fitness of mutants(*r*) and probability of going forward in each step (*λ*_*i*_), one can obtain the average number of steps needed for a single mutant to reproduce and increase the number of mutant to 1 *< k < N*.

By utilizing this method, a diagram depicting the number of mutants versus average time can be obtained for various structured populations. This diagram allows for capturing the evolutionary dynamics and understanding the progression towards fixation in populations with different structures. By examining the resulting diagrams, one can gain insights into the impact of population structure on the time it takes for mutants to fixate within a given population.

In reference [41], the challenging aspect of the problem lies in approximating the values of *λ*_*j*_. While for certain graph structures like complete graphs or cycle graphs, *λ*_*j*_ can be obtained accurately, for other complex structures such as Watts-Strogatz, Barabasi-Albert, and Erdos-Rnyi graphs, determining *λ*_*j*_ requires approximation methods. These approximation techniques are necessary due to the intricacies and randomness present in these types of graphs.

In this paper, the objective is to approximate the values of *λ*_*j*_ using machine learning methods in complex networks such as Watts-Strogatz, Barabasi-Albert, and Erdos-Rnyi graphs. Additionally, the proposed method is also applied to 2 and 3 dimensional periodic lattice network structures. The approach involves selecting appropriate features for training the machine learning model. To ensure the model’s accuracy and generalization, the machine learning algorithm is trained on graphs with a size smaller than 2000. However, the trained model is then applied to predict values for larger graph sizes, specifically 3000, 5000, and 10000. The simulation results and the predictions obtained from the machine learning method demonstrate agreement, indicating the effectiveness of the approach. This suggests that the machine learning method successfully captures the underlying patterns and dynamics of the Moran process in complex networks, allowing for accurate predictions even for larger graph sizes. Overall, this approach provides a valuable tool for approximating *λ*_*i*_ in complex networks and understanding the path to fixation in such systems.

## 2 General method

In this section, we will review the method introduced in reference [41] and explore how the application of machine learning for predicting *λ*_*j*_ can enhance the obtained results. The key metric for understanding the path of fixation in the Moran process is the average number of steps required for a single mutant to occupy *i* nodes of the network, denoted as *T*_*i*_. By obtaining the values of *T*_*i*_ for different numbers of occupied nodes, it is possible to construct a diagram illustrating the relationship between the number of mutants and the average time. This diagram provides specific insights into the trajectory and progression of fixation in the population. Following the methodology described in reference [41], the formula for calculating *T*_*i*_ can be expressed as follows:

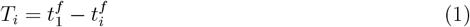

Where

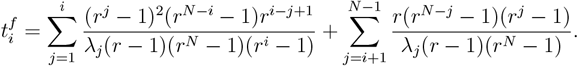

As mentioned earlier, with the knowledge of *λ*_*j*_ (the probability of the number of mutants increasing to *j* + 1), the size of the graph *N*, and the fitness parameter *r*, it is indeed possible to calculate *T*_*i*_. The precise formula for obtaining *T*_*i*_ depends on the specific approach or model for approximating *λ*_*j*_. However, in general, by combining the known values of *λ*_*j*_, *N*, and *r*, one can derive the average number of steps or time required for a single mutant to occupy *i* nodes of the network. In reference [41] coarse-graining methods applied for approximating *λ*_*j*_. Here we propose the use of machine learning methods for the same purpose.

Indeed, for the successful application of machine learning methods, it is important to select features that exhibit a non-zero correlation with *λ*_*j*_. By choosing relevant and informative features, we can ensure that the model captures the underlying patterns and dynamics of the Moran process accurately. As mentioned before for complete graph *λ*_*j*_ obtain without any approximation. The accurate value of *λ*_*j*_ for complete graph is

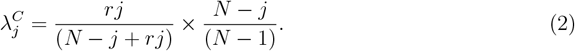

The first part of above relation is the probability that a mutant with fitness *r* chosen for reproduction. The second part of the equation represents the probability that, among the *N*−1 neighbors of the mutant, one of the *N*−*j* resident individuals is chosen to be replaced by the mutant’s offspring. By combining these two probabilities, the above equation provides 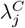 which represents the probability of the number of mutants increasing to *j* + 1 in a complete graph. For other complex network determining accurate value of *λ*_*j*_ is not possible. While the first part of 2 remains the same, representing the probability of a mutant being chosen for reproduction, the second part is influenced by the network structure. In contrast to a complete graph where all nodes are connected to each other, complex graphs exhibit a more intricate connectivity pattern where not all nodes are directly connected. As a result, the second part of 2 for complex networks will generally be smaller compared to that of a complete graph. This leads to a lower fixation rate for complex graphs compared to complete graph. For each complex graph, we introduce a new quantity denoted as *ω*_*j*_, defined as follows:

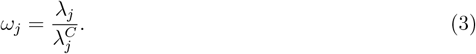

Since 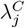 is larger than *λ*_*j*_ for all complex graphs and all values of *j*, the upper bound for *ω*_*j*_ is 1. In the process of solving a problem using machine learning methods, it is crucial to appropriately assign the target variable and features of the problem. Since *ω*_*j*_ scales between zero and one, it is a suitable quantity to be used as the target variable in the problem. By having *ω*_*j*_ and utilizing equation 3, the value of *λ*_*j*_ can be determined. In general *ω*_*j*_ depends on size of graph *N*, number of mutants *j*, fitness of mutants *r* and structured of graph. For complex graphs, two common features used in the problem are 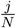 and *r*. In addition to these general features, other features that depend on the specific structures of the graph can be added on a case-by-case basis. These additional features can capture the characteristics of the network topology. By incorporating these features, which provide information about both the mutant population and the graph structure, into the machine learning model, we aim to improve the accuracy of predicting *λ*_*j*_ for complex networks.

## 3 Results and discussion

In this section, our focus is to examine the effect of network structure on the path to fixation in some complex networks. Our study will encompass the analysis of various complex network structures, including Erdos-Rnyi, Watts-Strogatz, and Barabasi-Albert graphs. Furthermore, to assess the effectiveness of our proposed method on regular graph structures, we will also evaluate 2 and 3 dimensional periodic lattice network configurations. In all cases, we employed machine learning techniques to train a model. The target variable in our training process was defined as *ω*_*j*_, which represents the desired output or prediction. We selected relevant features to input into the machine learning model, including the proportion of mutants in the population 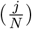, the fitness parameter (*r*), and other variables that are derived from the structure of the specific graph under consideration. These additional variables, which depend on the network structure, are chosen based on their potential correlation with the target variable *ω*_*j*_. By incorporating these features, we aim to capture the relevant information and characteristics of the graph that can enhance the accuracy and effectiveness of our machine learning model. Additionally, for each specific case, we employed different machine learning models that yielded the best results. For Erdos-Rnyi, Watts-Strogatz, and Barabasi-Albert graphs, we utilized a linear regression model and set the polynomial degree of the features to two. This choice of model and feature transformation was found to be most effective in capturing the relationships between the features and the target variable in these network structures. On the other hand, for 2 and 3 dimensional periodic lattice networks, we opted for a random forest regressor as the machine learning model. Similarly, we set the polynomial degree of the features to two to account for potential non-linear relationships. This choice of model was found to be more suitable for capturing the complex dynamics and patterns present in regular lattice network structures. Our training test generated among graph with size smaller than 2000 while we compare results obtain from machine learning with simulation results in graph with size 3000,5000, and 10000. The agreement observed between the machine learning predictions and the simulation results for these large graphs is a positive indication of the success of the machine learning approach. In all the cases investigated in this study, we utilized the scikit-learn Python library to apply machine learning methods and NetworkX python library for generating graphs.

### 3.1 Erdos-Rnyi graph

In Erdos-Rnyi graph, every pair of nodes is connected to each other with a certain probability, denoted as *p*. Most structure properties of Erdos-Rnyi graph summarized in value of *p*. It appears that apart from 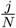 and *r, p* is also a relevant feature for predicting *ω*_*j*_ in the Erdos-Rnyi graph. We used a linear regression model and set the polynomial degree of the features to two. Figure 1 shows the path of fixation for Erdos-Rnyi graph with size 3000,5000 and 10000 and *p* = .1.

**Fig 1.**
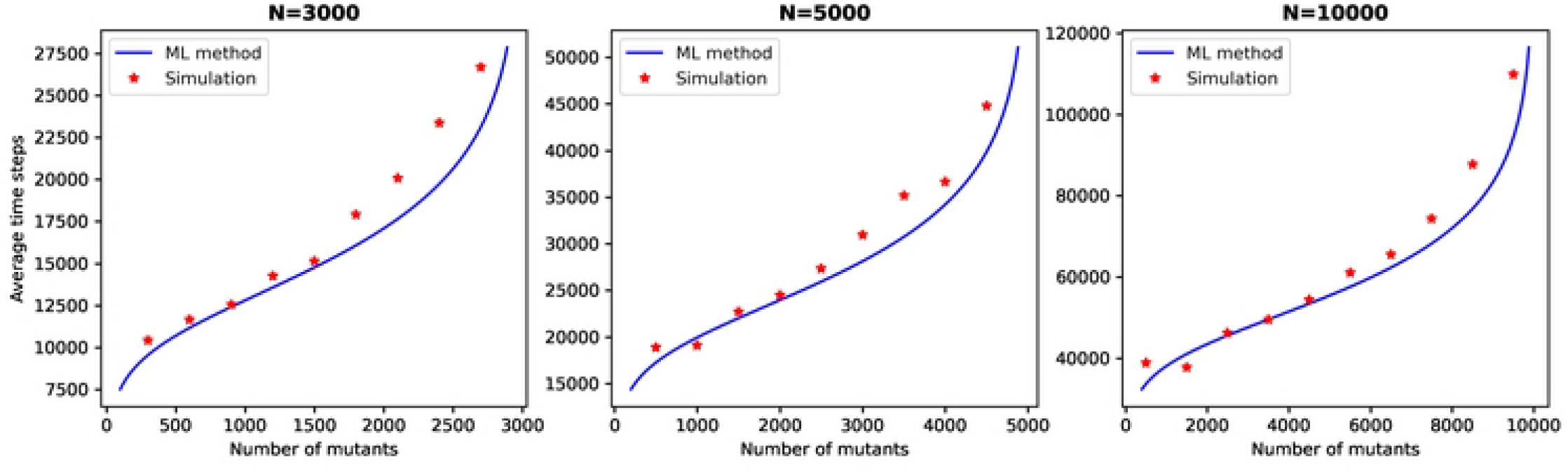

### 3.2 Watts-Strogatz graph

To generate a Watts-Strogatz graph, we begin with a regular graph which each node has 2*m* edges and then, by rewiring probability *P*_*WS*_, rewire the edges. This procedure, with proper choices for *P*_*WS*_, leads to a graph with small-world property. In fact parameter *P*_*WS*_ controls the level of randomness or regularity in the graph structure. Based on our analysis in Watts-Strogatz graph, besides 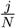 and *r, P*_*WS*_ and 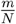 are good features for predicting *ω*_*j*_. In trained set we used graph with size smaller than 2000 and 0.01 *< P*_*WS*_ *<* .99. Figure 2 shows the path of fixation for Watts-Strogatz graph with size 3000,5000 and 10000 and *P*_*WS*_ = .1.

**Fig 2.**
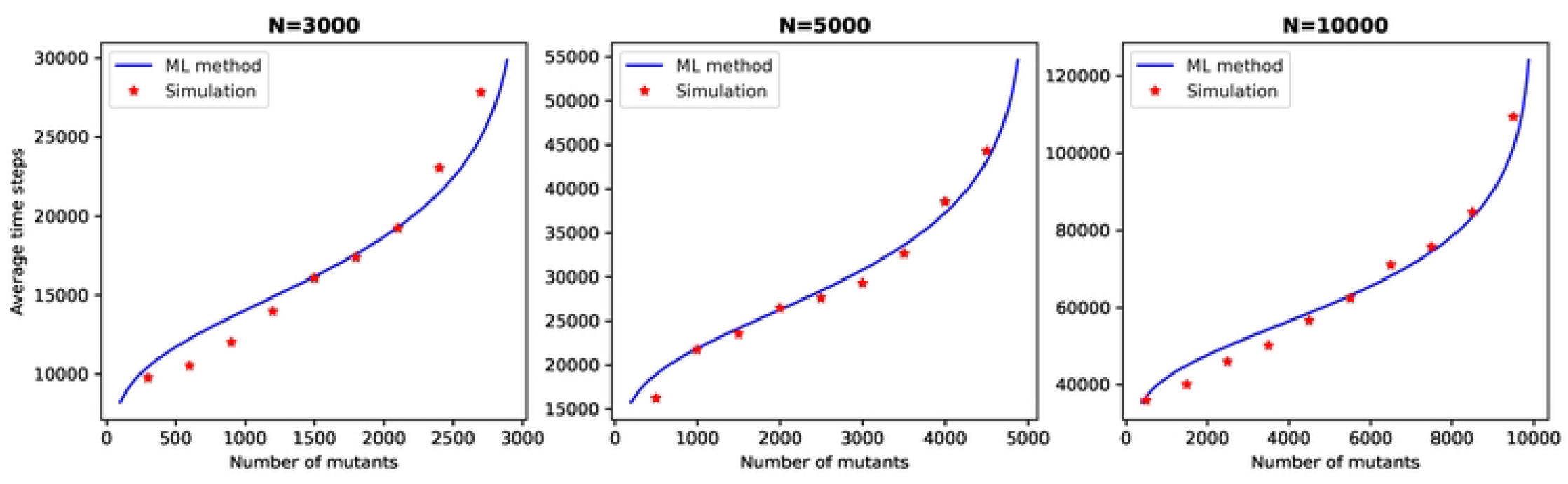

### 3.3 Barabasi-Albert graphs

Barabasi-Albert graphs, also known as scale-free graphs, are constructed by starting with a complete graph of size *m*_0_. Nodes are then incrementally added to the graph, and each added node connects to *m* existing nodes with a probability proportional to the degree of the existing nodes. This process continues until the desired network size is achieved. According to our study, neither *m* nor *m*_0_ are deemed suitable as features to be considered in the machine learning process. Best results for coinciding simulation results and machine learning results obtain when features of problem are 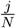 and *r*. The degree distribution of a scale-free graph follows a power-law distribution regardless of the values of *m* and *m*_0_. Therefore, it is not surprising that *m* and *m*_0_ do not play a critical role in the learning process.

Figure 3 shows the path of fixation for Barabasi-Albetr graph with size 3000,5000 and 10000.

**Fig 3.**
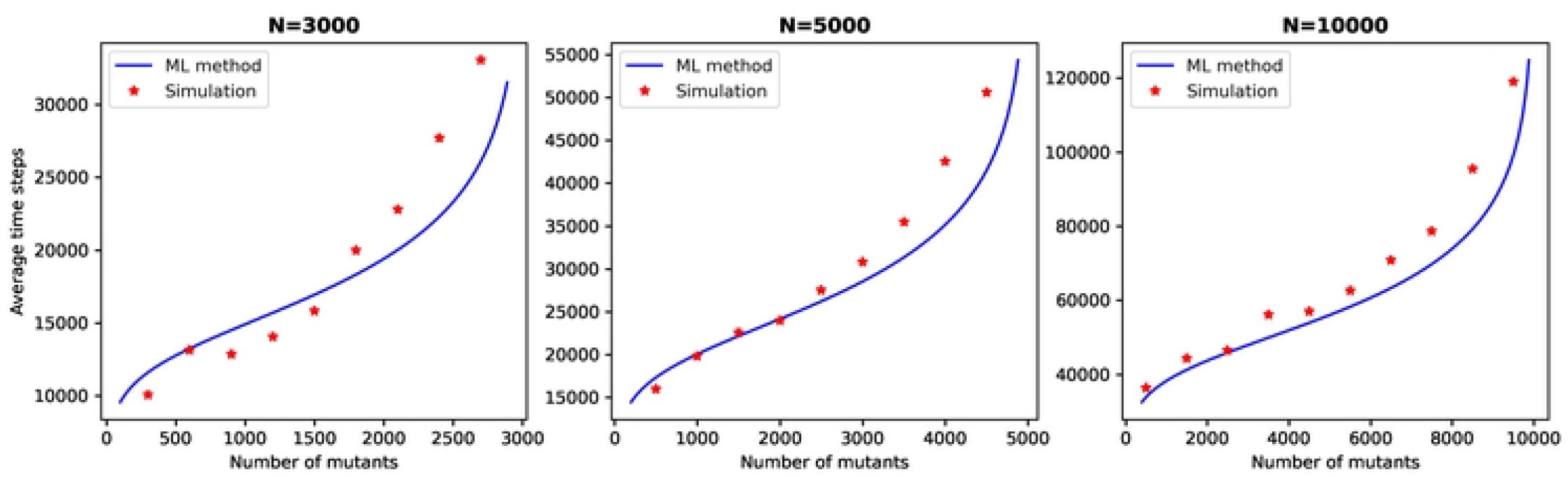

### 3.4 2 and 3 dimensional periodic lattice graph

To investigate the effectiveness of our method on regular graphs, we applied a machine learning approach to 2-dimensional and 3-dimensional periodic lattice graphs. The training of the model involved graphs with a size smaller than 2000. The features used in the model were 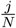 and *r*, and a random forest regressor was employed with a polynomial degree set to two.

The figure 4 presented depict the simulation and machine learning results for the 2-dimensional periodic lattice graphs with sizes of 3600, 4900, and 10000. Additionally, the figure 5 display the same for the 3-dimensional periodic lattice graphs with sizes of 3375, 4913, and 10648. It is worth noting that the agreement between the simulation and machine learning results is deemed satisfactory.

**Fig 4.**
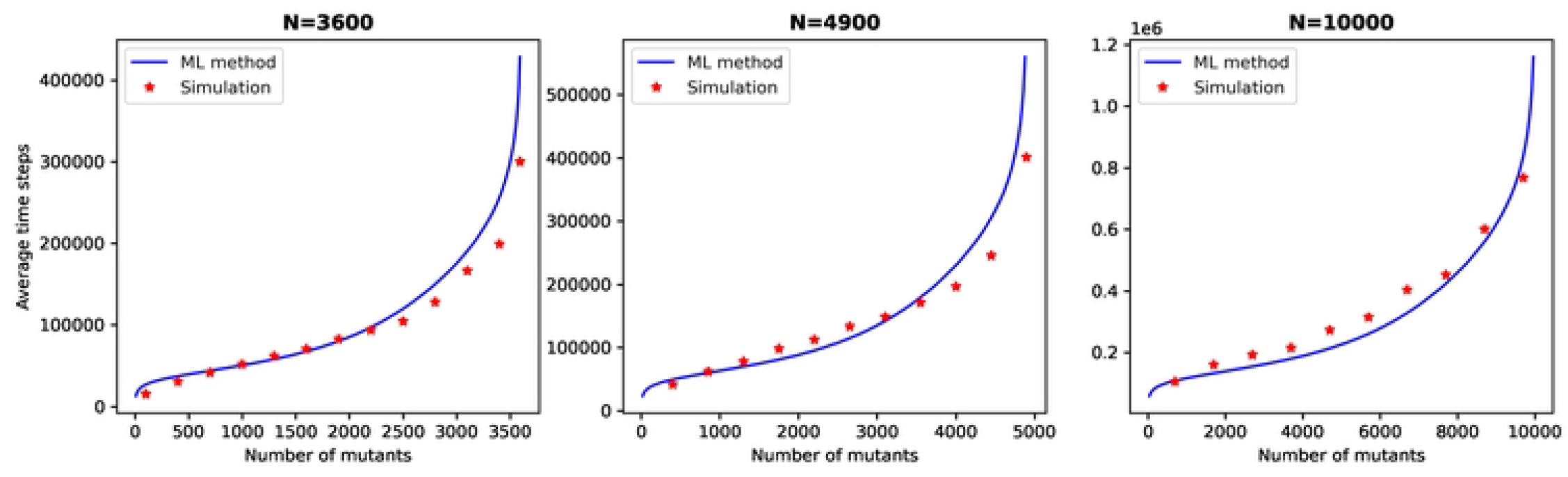

**Fig 5.**
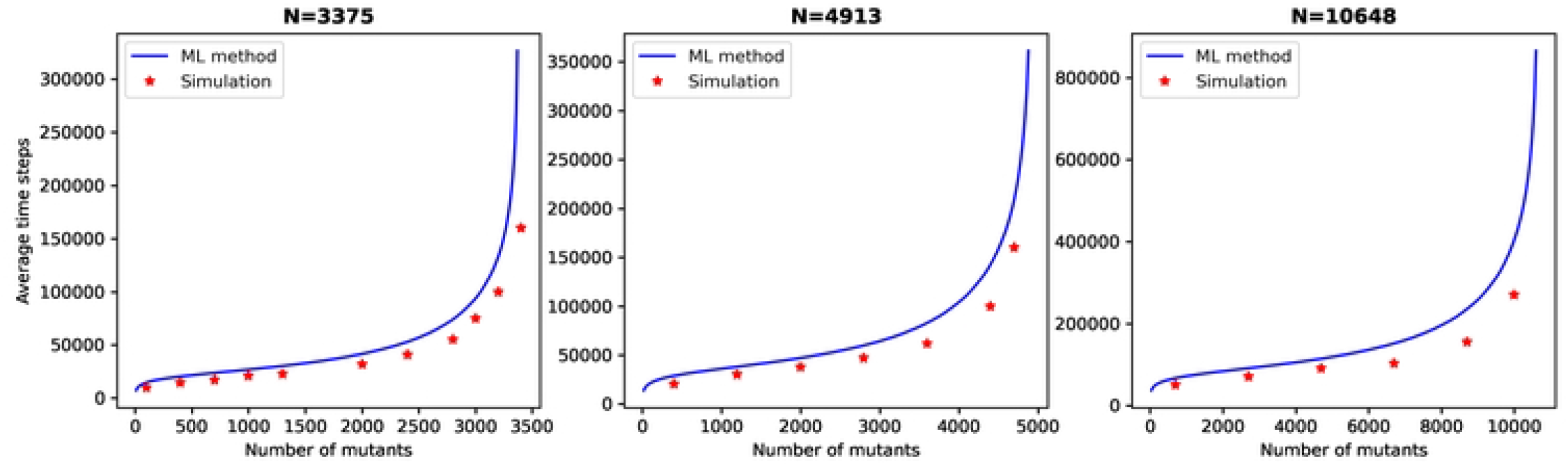

## 4 Summary and concluding remarks

In this paper, our focus is on exploring the predictive capabilities of machine learning methods in determining the path of fixation in Moran processes within various structured populations. Our objective was not to directly predict the path of fixation. Instead, we aimed to utilize machine learning methods to predict *λ*_*j*_, which represents the probability of the number of mutants increasing to *j* + 1 when there are *j* mutants in the population within the Moran process. Based on the previous work conducted in analyzing the Moran process, it is indeed possible to identify the path to fixation by just having the values of *λ*_*j*_ for all 1 *< j < N*. These values provide information about the probabilities of the number of mutants increasing to different population sizes as the process evolves. By utilizing this data, one can infer and determine the path leading to fixation within the Moran process. The application of machine learning methods has been successful in both complex and regular graphs. Since simulation of Moran process through large graph needs significant time and computational resources, the utilization of machine learning methods can effectively save both time and computational facilities. By training a machine learning model using small and medium-sized graphs, one can leverage the predictive capabilities of the model to make predictions for larger graphs, thus reducing the need for extensive simulations. This approach offers a more efficient and resource-saving alternative for analyzing and understanding complex networks. This approach could be highly useful in analyzing diffusion-like processes in meta-populations [42–44]. In meta-populations, it is possible to train a machine learning model using several sub-graphs of the population. By leveraging the predictive capabilities of the model, one can then use it to determine the path of the diffusion process across the entire population.

Code related to this article is hosted on

https://github.com/mehdiphy/Machine-learing-method-in-Moran-process

